# A robust human norovirus replication model in zebrafish larvae

**DOI:** 10.1101/528364

**Authors:** Jana Van Dycke, Annelii Ny, Nádia Conceição-Neto, Jan Maes, Myra Hosmillo, Arno Cuvry, Ian Goodfellow, Tatiane C. Nogueira, Erik Verbeken, Jelle Matthijnssens, Peter de Witte, Johan Neyts, Joana Rocha-Pereira

**Author notes:** Janssen Pharmaceutical Companies of Johnson and Johnson, Beerse, Belgium (Nádia Conceição-Neto).

## Abstract

Human noroviruses (HuNoVs) are an important cause of epidemic and endemic acute gastroenteritis worldwide; annually about 700 million people develop a HuNoV infection resulting in ∼219,000 deaths and a societal cost estimated at 60 billion US dollars ^1^. The lack of robust small animal models has significantly hindered the understanding of norovirus biology and the development of effective therapeutics against HuNoV. Here we report that HuNoV GI and GII replicate to high titers in zebrafish (*Danio rerio*) larvae; replication peaks at day 2 post infection and is detectable for at least 6 days. HuNoV is detected in cells of the hematopoietic lineage, the intestine, liver and pancreas. Antiviral treatment reduces HuNoV replication by >2 log_10_, showing that this model is suited for antiviral studies. Downregulation of *fucosyltransferase 8 (fut8)* in the larvae reduces HuNoV replication, highlighting a common feature with infection in humans. Zebrafish larvae constitute a simple and robust replication model that will largely facilitate studies of HuNoV biology and the development of antiviral strategies.

---

Large outbreaks of norovirus gastroenteritis are frequent and have a significant impact in terms of morbidity, mortality and health care costs, in particular in hospital wards and nursing homes. Chronic norovirus infections present a problem for a large group of immunodeficient patients, who may present with diarrhea for several months. Furthermore, in countries where routine rotavirus vaccination has been implemented, noroviruses are the most common cause of severe childhood diarrhea resulting in important morbidity and mortality ^2^. Knowledge on the biology and pathogenesis of human noroviruses largely depends upon the development of robust and physiologically relevant cultivation systems. A number of such model systems have been reported in recent years but still carry important limitations. HuNoV replication has been reported in large animals such as chimpanzees, gnotobiotic pigs and calves. However these animals are either not suited for extensive studies or are, in the case of chimpanzees no longer allowed due to ethical reasons ^3-5^. Importantly, a HuNoV mouse model was described in BALB/c Rag-γ c-deficient mice, but only a short-lasting replication was achieved, which limits its applications^6^. Standard cell culture models are to date not available, but first steps towards this have been given by establishing that (i) human B-cells are susceptible to HuNoV and that (ii) HuNoV can be cultivated in stem-cell-derived enteroids. ^7-9^ There is thus an urgent need for simpler, more robust, widely available HuNoV replication models. Such models should contribute to a better understanding of the biology of HuNoV replication and infection, this will significantly facilitate larger-scale research efforts, such as the development of therapeutic strategies.

Zebrafish *(Danio rerio)* are optically-transparent tropical freshwater fish of the family *Cyprinidae* that are widely used as vertebrate models of disease. They have remarkable genetic, physiologic and pharmacologic similarities to humans. Compared to rodents, the maintenance and husbandry costs are very low. Zebrafish have high fecundity and using their offspring is in better compliance with the 3Rs principles of humane animal experimentation (EU Directive 2010/63/EU). The immune system of zebrafish is comparable to that of humans; there are B and T cells, macrophages, neutrophils and a comparable set of signaling molecules and pathways ^10^. Whereas innate immunity is present at all developmental stages, adaptive immunity develops after 4-6 weeks of life ^11,12^ Host-pathogen interactions can be studied, as zebrafish are naturally infected by multiple bacteria, protozoa and viruses that affect mammals ^11^. Infection of zebrafish larvae has been shown with various pathogens including *Mycobacterium* ^13^, some human viruses (herpes simplex 1, influenza A, hepatitis C and chikungunya viruses) ^14-17^ and enteric bacteria, e.g. *E. coli, Listeria, Salmonella, Shigella* and *Vibrio* ^11^. The intestinal tract of zebrafish is comprised of large folds of an epithelial lining, a *lamina propria* containing immune cells and underlying smooth muscle layers ^18^. Enterocytes, goblet cells, enteroendocrine cells, and possibly M-cells are present, but not Paneth cells or Peyer’s patches ^12^. Intestinal tuft cells, which were recently shown to be a target cell for the mouse norovirus (MNV) ^19^, have been described in teleost fish thus are likely present in zebrafish. There is an intestinal bulb (instead of a stomach) and a mid-and posterior intestine ^18^. Epithelial cells show a high-turnover from base to tip, with intestinal epithelial stem cells at the base and apoptotic cells at the tips ^18^. A resident commensal microbiota is present (comprising most bacterial phyla of mammals) and serve analogous functions in the digestive tract ^11,18^. Here we report a robust replication model of HuNoV in zebrafish larvae.

Zebrafish larvae were infected with a PBS suspension of a HuNoV positive stool sample at 3 days post-fertilization (dpf). At this time point, zebrafish larvae have hatched and organs are formed (including the full length gastrointestinal tract). Three nL, containing 3.4×10^6^ viral RNA copies of HuNoV GII.P7-GII.6 (1.1×10^13^ RNA copies/g of stool), were injected in the yolk of the larvae (which provides nutrition during early larval stage). Each day post infection (pi), the general condition of the zebrafish larvae was assessed microscopically and these were harvested in groups of 10 for viral RNA quantification by RT-qPCR^16^. To detect input virus, in every independent experiment, 10 larvae were harvested at day 0 pi (specifically 1 h pi). A maximum increase of ∼2.5 log_10_ in viral RNA copies compared to day 0 was detected at day 2 pi (Fig. 1A); high levels of viral RNA remained detectable for at least 6 days pi (Fig. 1A, S1). When larvae were injected with 3 nL of UV-inactivated HuNoV GII.P7-GII.6, no increase in viral RNA titers was detected (Fig. 1A). No obvious signs of distress or disease were observed as a result of the infection (e.g. changes in posture, swimming behavior or signs of edema). To determine the 50% infectious dose (ID_50_) for this strain, larvae were infected with 10-fold dilution series of the virus (Fig. 1B). The ID_50_ was calculated to be 1.8 × 10^3^ viral RNA copies. HuNoV antigens were detected using the commercial enzyme immunoassay (EIA) RIDASCREEN (R-Biopharm), in HuNoV GII.P7-GII.6-infected zebrafish larvae harvested at day 3 pi (Fig. 1C). We next investigated the innate immune response of 3 dpf larvae to a HuNoV infection. An increased expression of *ifn, mx* and *rsad2/viperin* mRNA was detected, with a 7-fold, 23-fold and 105-fold maximum increase, respectively, when compared to control inoculated larvae (Fig. 1D). This level of upregulation is in line with the observed induction of the IFN response following an influenza A infection of zebrafish larvae ^16^. These same genes (or the related cytokines) were detected in other *in vivo* models, such as in HuNoV-infected calves or MNV-infected mice ^5,20^, or in a HuNoV replicon system in the case of viperin ^21^. All together this points out that the antiviral signaling cascades which are activated upon a HuNoV infection of zebrafish larvae are relevant and likely the same as in humans. Zebrafish are thus a suitable model for the study of the innate immune response to a HuNoV infection.

**Fig. 1.**
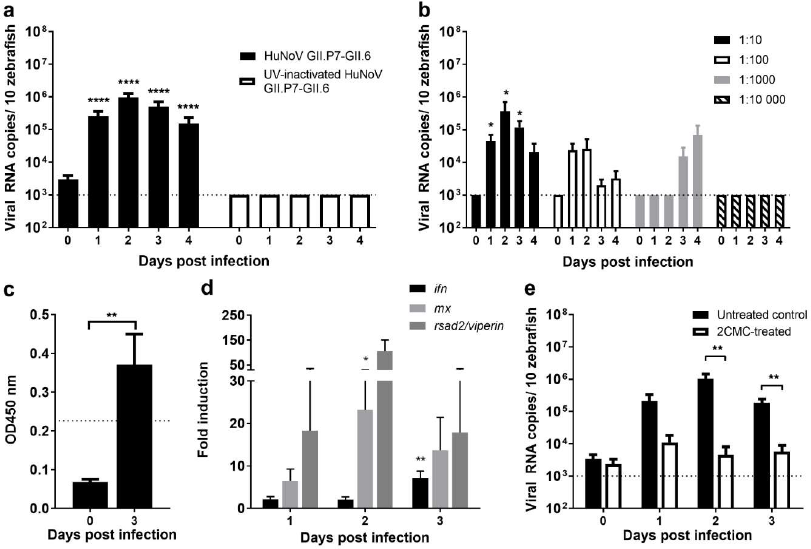
HuNoV GII.P7-GII.6 replicates in zebrafish larvae. **(a)** Infection of zebrafish larvae (3dpf) with HuNoV GII.P7-GII.6 (22 independent experiments) or UV-inactivated virus (2 independent experiments). Bars represent viral RNA levels/10 larvae (per condition in each experiment), quantified by RT-qPCR. The dotted line represents the limit of detection (LOD). Values of viral RNA in larvae infected with the UV-inactivated sample was set at the LOD (undetected by RT-qPCR). **(b)** Zebrafish larvae were injected with serial dilutions (ranging from 1:10-1:10,000) of HuNoV GII.P7-GII.6 (5 independent experiments). Zebrafish larvae were harvested each day pi, bars represent the mean values ± SEM of viral RNA levels/10 zebrafish larvae as quantified by RT-qPCR. The dotted line represents the LOD. **(c)** Viral antigens were quantified by ELISA in HuNoV GII.P7-GII.6-infected larvae. Bars represent OD values/10 larvae. The dotted line is the calculated cutoff+10%, above which samples are considered positive. **(d)** The effect of a HuNoV GII.P7-GII.6 infection on the expression of *ifn, mx* and *rsad2/viperin* mRNA was determined by RT-qPCR, normalized to β-actin (9 independent experiments). Bars represent the fold-induction of *ifn, mx* and *rsad2/viperin* mRNA in HuNoV-infected larvae/10 larvae. **(e)** HuNoV GII.P7-GII.6-infected larvae were treated with 4 mM of 2’-C-methylcytidine via immersion in Danieau’s solution. Treatment started one day before infection and was refreshed every 12 hours (4 independent experiments). Bars represent the viral RNA levels/10 zebrafish larvae. For all graphs: in every independent experiment 10 zebrafish larvae were harvested at each time point, mean values ± SEM are presented, Mann-Whitney test, where ****p<0.0001, **p<0.01, *p<0.05.

Next, infected larvae were treated with a broad-spectrum antiviral, i.e. the viral polymerase inhibitor 2’-C-methylcytidine (2CMC) of which we showed earlier inhibition of MNV replication *in vitro* and in mice ^22,23^, by immersion (whereby the molecule was added to the water). A 2.4 log_10_ reduction in viral RNA titers was observed at the peak of replication (Fig. 1E). The fact that replication can be significantly reduced with an inhibitor of the viral polymerase, provides further evidence that HuNoV replicates efficiently in zebrafish larvae and that the model is suitable for antiviral drug development.

One of the hurdles to develop robust replication models for HuNoV is the need to use a stool sample of an infected patient as inoculum. In order to rule out any potential impact of other agents present in the sample we fully characterized the samples used. A viral metagenomics analysis was performed on the clinical sample containing the HuNoV GII.P7-GII.6 used in this study, together with subsequent clinical samples of the same chronically-infected 2.5-year old transplant patient (Fig. S2). The viral population consisted predominantly of HuNoV (with a minor presence of anelloviruses, common in patients undergoing immunosuppressive therapy ^24^). Mutations that occurred in the virus over the course of the infection were mostly in the capsid-encoding region (Fig. S2, Table S1) and did not affect the kinetics of virus replication in larvae (Fig. S2B-D). Infection of zebrafish larvae with other HuNoV genotypes was next performed. Infection with the HuNoV GII.P4 New Orleans-GII.4 Sydney strain, recovered from stool samples of two different patients, resulted in a >3 log_10_ increase in viral replication in both cases. A maximum of ∼10^7^ viral RNA copies/10 zebrafish larvae was detected at day 2 pi (Fig. 2A), the highest observed in this model ^6,8,9^. Viral non-structural and structural antigens were detected by western blot (Fig. 2B, Fig. S3) and by EIA (Fig. 2C), respectively. Viral antigens were no longer detected by EIA in 2CMC-treated HuNoV GII.4-infected larvae (Fig. 2C). Infection with HuNoV GII.P16-GII.2 (Fig. 2D) and GII.P16-GII.3 (Fig. 2E) yielded increasing viral RNA titers, although the replication kinetics of GII.P16-GII.3 was slower than that observed for the other genotypes. Slower kinetics of HuNoV GII.3 replication was also observed in stem-cell-derived enteroids ^9^. A GI HuNoV, specifically GI.P7-GI.7, replicated with comparable kinetics to GII viruses (Fig. 2F).

**Fig. 2.**
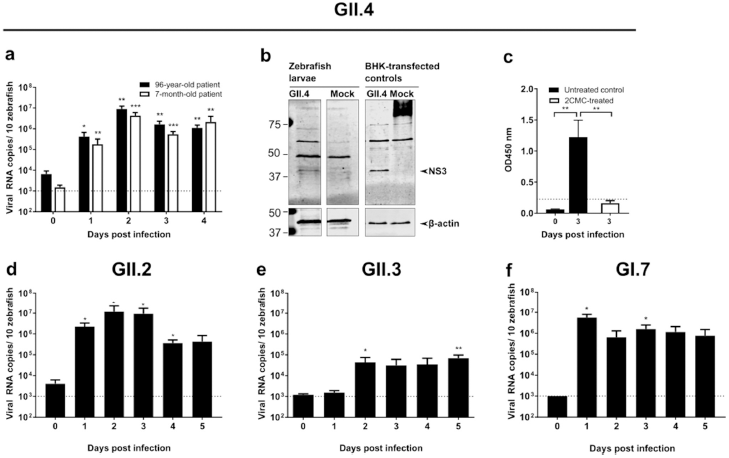
Infection of zebrafish larvae with HuNoV GI and GII of other genotypes. **(a)** Zebrafish larvae infected with GII.P4 New Orleans-GII.4 Sydney, from a 96-year-old patient (black bars) and a 7-month-old patient (empty bars), were harvested each day pi (6 independent experiments). Bars represent viral RNA levels/10 zebrafish larvae, quantified by RT-qPCR. The dotted line represents the limit of detection (LOD). **(b)** The viral NS3 protein was detected in HuNoV GII.4-infected larvae by western blot analysis, BHK cells transfected with a GII.4 construct were used as positive control. **(c)** Structural antigens were detected by EIA [here 2CMC-treated HuNoV GII.4-infected zebrafish were also included]. Bars represent OD values/10 zebrafish larvae. The dotted line is the calculated cutoff+10%, above which samples are considered positive. **(d)** Zebrafish larvae infected with HuNoV GII.P16-GII.2 (from an 87-year-old patient) were harvested each day pi (5 independent experiments). Bars represent the viral RNA levels/10 zebrafish larvae as quantified by RT-qPCR. (E) Zebrafish larvae infected with GII.P16-GII.3 (from a 3.5-year-old patient) were harvested each day pi (5 independent experiments). Bars represent the viral RNA levels/10 zebrafish larvae as quantified by RT-qPCR. **(f)** Zebrafish larvae infected with HuNoV GI.7 (from a 52-year-old patient) were harvested each day pi (7 independent experiments). Bars represent the viral RNA levels/10 zebrafish larvae as quantified by RT-qPCR. The dotted line represents the LOD. In all graphs: in every independent experiment 10 larvae were harvested at each time point, mean values ± SEM are presented, Mann-Whitney test, where ***p<0.001, **p<0.01, *p<0.05.

Infection of zebrafish larvae with MNV (genogroup V) yielded no productive infection (Fig. S4), most likely due to the fact that the receptors CD300lf and CD300ld are not encoded by zebrafish ^25^. Since HuNoV infections typically occur *via* the oral route, we next attempted to infect larvae *via* immersion. No consistent increase in viral replication was noted up to day 5 pi (data not shown). To determine the preferential site of replication of HuNoV in zebrafish larvae, HuNoV GII.P7-GII.6-infected larvae were dissected at day 3 pi in 4 different parts (yolk, head, body and tail). Viral RNA titers were detected in every part (Fig. 3A), whereby the yolk (the initial site of inoculation) had the lowest titers, implying that HuNoV disseminates past the yolk and intestine. To investigate which tissues are infected, sagittal and coronal histological sections of larvae infected with HuNoV GII.P7-GII.6 were stained with HuNoV VP1-specific antibodies (Fig. 3B, Fig. S5-6). Viral antigens were frequently detected in the intestinal bulb, pancreas and liver. A strong signal was observed in the caudal hematopoietic tissue (CHT), which contains hematopoietic stem/progenitor cells (HSPCs) that differentiate into multiple blood lineages and by 4 dpf start to migrate to the kidney marrow and thymus ^26^. This migration may explain why HuNoV was detected in every section of the larvae. HuNoV has been detected in the intestine and liver of chimpanzees ^3^, and has as well been reported to replicate in cells of hematopoietic lineage ^27^.

**Fig. 3.**
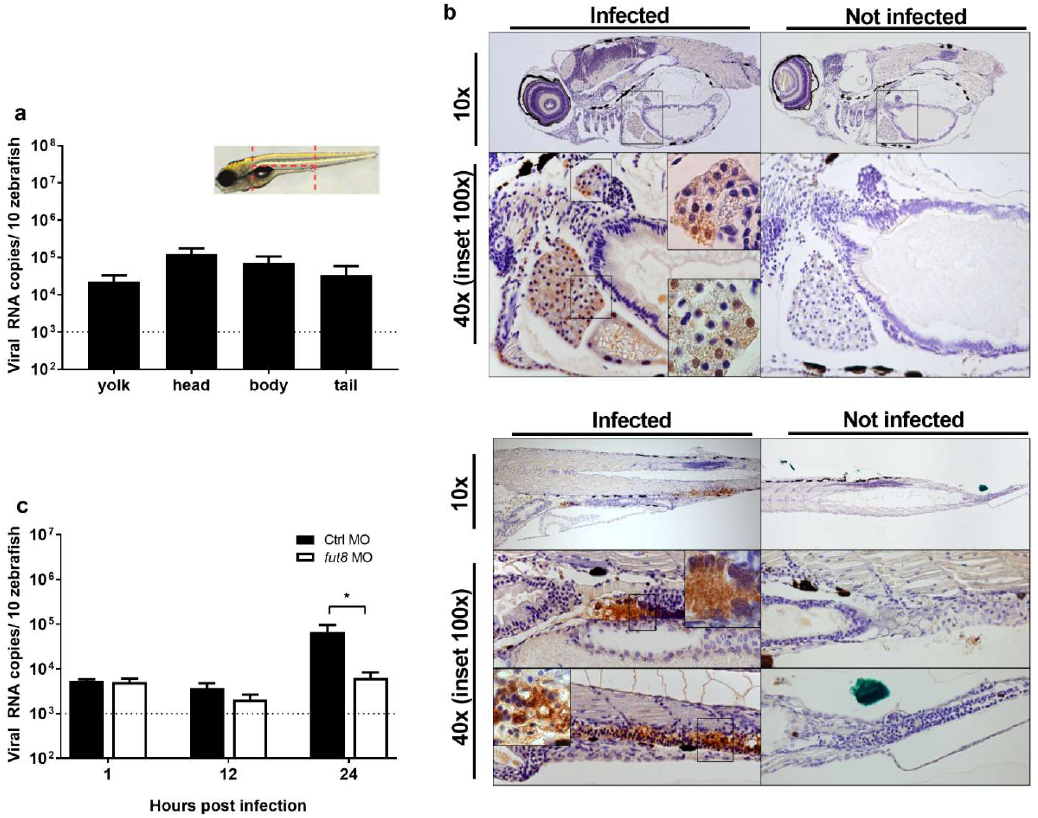
HuNoV sites of replication and relevance of a *fucosyltransferase* gene in susceptibility of zebrafish larvae. **(a)** Ten HuNoV-infected larvae were deyolked and then dissected into head, body, and tail (as depicted in the scheme). Bars represent the mean values ± SEM of viral RNA levels as quantified by RT-qPCR (7 independent experiments) **(b)** Immunohistochemistry of sagittal sections of HuNoV GII.P7-GII.6-infected zebrafish larvae harvested at day 3 pi (and the respective uninfected controls) was performed. Images show 5 μm sections stained with antibodies targeting VP1 at 10x and 40x magnifications, insets at 100x magnification. In the top panel, viral antigens were detected in the liver and pancreas; the lower panel depicts staining in the intestine and CHT of infected zebrafish larvae **(c)** One-cell stage embryos were microinjected with a control morpholino or a morpholino targeting *fut8,* and subsequently infected at 3 dpf with HuNoV GII.P7-GII.6. Ten zebrafish larvae per independent experiment were harvested at 1, 12 and 24 h pi. Bars represent the mean values ± SEM of viral RNA levels as quantified by RT-qPCR (6 independent experiments). In all graphs: The dotted line represents the LOD, Mann-Whitney test, where *p<0.05.|

The HuNoV capsid binds to fucose-containing histo-blood group antigens (HBGAs), which are key players in determining susceptibility to infection ^28^. Fucosyltransferase *(FUT)* 2 and 3 genes determine the addition of terminal fucose residues to HBGAs; individuals with a null *FUT2* allele are known to be resistant to infection with many HuNoV strains ^29^. Zebrafish have *fut7-ll* genes, of which the *fUt8* gene has the highest similarity between human, mice and zebrafish, and is the only fUt gene that is expressed in tissues that are relevant to a norovirus infection (Table S2). Fut8 is the single enzyme responsible for adding core fucose (a-1,6-Fuc) both in zebrafish and mammals. When *fUt8* expression was knocked-down by injection of *fut8* targeting morpholino (MO) antisense oligonucleotides (Fig. S7), lower levels of HuNoV and a slower kinetics of replication was observed, as compared to those injected with a control MO (Fig. 3). This demonstrates that *fut8* is involved in susceptibility of zebrafish to HuNoV. We hypothesize that core fucosylation plays a regulatory role in HuNoV susceptibility by either regulating the binding to glycoproteins and/or their expression levels at the cell surface, as was also reported for the hepatitis B virus ^30^.

In conclusion, here we describe a robust and very convenient HuNoV replication model in zebrafish larvae. While zebrafish share about ∼70% of their genes with humans, there are obvious differences with the natural host. However, it is a whole-organism with complete organs and systems, thus providing a unique chance to identify physiologically relevant features of a HuNoV infection. In addition, the zebrafish model brings unique advantages, their optical transparency allows live imaging studies, for example using transgenic lines with tagged immune or gut cells, which would aid studies of tissue tropism and pathogenesis. The availability of simple genetic manipulation methods facilitates the understanding of gene functions, which, combined with the availability of many knockout alleles ^31^, could significantly enhance our ability to dissect HuNoV-host interactions. Zebrafish are widely available at universities/research centers and their use is amenable to high-throughput studies. A trained researcher can inject/manipulate hundreds of larvae per day requiring only a microscope, a micromanipulator and injection pump. Moreover, only minute amounts of virus (few nL) are required to infect zebrafish larvae. Consequently, a 100 mg stool aliquot (with a high virus titer) is sufficient to inject about 300,000 larvae, yielding 30,000+ data points. The ability to generate large and homogenous datasets is essential for large-scale efforts such as the development of therapeutics. This model, here validated for antiviral drug studies using GI and GII HuNoVs, now allows to readily assess the potential antiviral activity of novel inhibitors. Overall this model is a major step forward in the study of HuNoV replication and provides the first robust small laboratory animal of HuNoV infection.

## Acknowledgments

We very much appreciate the expert technical assistance and dedication of Jasper Rymenants, Lindsey Bervoets, Charlotte Vanderheydt, Kathleen Van den Eynde, Wilfried Versin and value the expert advice of Prof. Lieve Moons, Dr. Jessie Van houcke and Annelies Van Dyck (Lab. Animal Physiology and Neurobiology, KU Leuven). We thank Dr. Peter Sander (R-Biopharm) for kindly providing antibodies and the EIA kit. We thank the CEMOL Molecular Diagnostic department of the University Hospital of Leuven for the collaboration. JRP and the research leading to these results has received funding from the People Programme (Marie Curie Actions) of the European Union’s Seventh Framework Programme (FP7/2007-2013) under REA grant agreement n° 608765. JVD is an SB doctoral fellow of the Scientific Fund for Research of Flanders (FWO) and NCN is an SB doctoral fellow of Flanders Innovation and Entrepreneurship (VLAIO). IG and MH are supported by funding from the Wellcome Trust (Refs: 097997/Z/11/Z and 207498/Z/17/Z). The authors declare no conflicts of interest.

## Author contributions

JVD, AN, and JRP designed the experiments. JVD, AN, NCN, JM, MH and AC performed experiments, EV supervised the immunohistochemistry experiments. JVD, NCN and JRP analyzed the results. JVD and JRP wrote the manuscript. IG, TCN, JMT, PdW, JN and JRP were involved in designing the concept and planning and supervising the work. All authors discussed the results and contributed to the final manuscript.

## Supplementary Materials

Materials and Methods

Figures S1-S7

Tables S1-S4

References (31-46)

## References

1 Bartsch, S. M., Lopman, B. A., Ozawa, S., Hall, A. J. & Lee, B. Y. Global Economic Burden of Norovirus Gastroenteritis. PloS one 11, e0151219, doi:10.1371/journal. pone.0151219 (2016).

2 Hemming, M. et al. Major reduction of rotavirus, but not norovirus, gastroenteritis in children seen in hospital after the introduction of RotaTeq vaccine into the National Immunization Programme in Finland. European journal of pediatrics 172, 739–746, doi:10.1007/s00431-013-1945-3 (2013).

3 Bok, K. et al. Chimpanzees as an animal model for human norovirus infection and vaccine development. Proc Natl Acad Sci U S A 108, 325–330, doi:10.1073/pnas.1014577107 (2011).

4 Cheetham, S. et al. Pathogenesis of a genogroup II human norovirus in gnotobiotic pigs. J Virol 80, 10372–10381, doi:10.1128/JVI.00809-06 (2006).

5 Souza, M., Azevedo, M. S., Jung, K., Cheetham, S. & Saif, L. J. Pathogenesis and immune responses in gnotobiotic calves after infection with the genogroup II.4-HS66 strain of human norovirus. J Virol 82, 1777–1786, doi:10.1128/JVI.01347-07 (2008).

6 Taube, S. et al. A mouse model for human norovirus. MBio 4, e00450–00413, doi:10.1128/mBio.00450-13 (2013).

7 Jones, M. K. et al. Human norovirus culture in B cells. Nat Protoc 10, 1939–1947, doi:10.1038/nprot.2015.121 (2015).

8 Jones, M. K. et al. Enteric bacteria promote human and mouse norovirus infection of B cells. Science 346, 755–759, doi:10.1126/science.1257147 (2014).

9 Ettayebi, K. et al. Replication of human noroviruses in stem cell-derived human enteroids. Science 353, 1387–1393, doi:10.1126/science.aaf5211 (2016).

10 Goody, M. F., Sullivan, C. & Kim, C. H. Studying the immune response to human viral infections using zebrafish. Developmental and comparative immunology 46, 84–95, doi:10.1016/j.dci.2014.03.025 (2014).

11 Kanther, M. & Rawls, J. F. Host-microbe interactions in the developing zebrafish. Current opinion in immunology 22, 10–19, doi:10.1016/j.coi.2010.01.006 (2010).

12 Lewis, K. L., Del Cid, N. & Traver, D. Perspectives on antigen presenting cells in zebrafish. Developmental and comparative immunology 46, 63–73, doi:10.1016/j.dci.2014.03.010 (2014).

13 Ramakrishnan, L. The zebrafish guide to tuberculosis immunity and treatment. Cold Spring Harb Symp Quant Biol 78, 179–192, doi:10.1101/sqb.2013.78.023283 (2013).

14 Burgos, J. S., Ripoll-Gomez, J., Alfaro, J. M., Sastre, I. & Valdivieso, F. Zebrafish as a new model for herpes simplex virus type 1 infection. Zebrafish 5, 323–333, doi:10.1089/zeb.2008.0552 (2008).

15 Ding, C.-B. et al. Zebrafish as a Potential Model Organism for Drug Test Against Hepatitis C Virus. PloS one 6, e22921, doi:10.1371/journal.pone.0022921 (2011).

16 Gabor, K. A. et al. Influenza A virus infection in zebrafish recapitulates mammalian infection and sensitivity to anti-influenza drug treatment. Dis Model Mech 7, 1227–1237, doi:10.1242/dmm.014746 (2014).

17 Palha, N. et al. Real-time whole-body visualization of Chikungunya Virus infection and host interferon response in zebrafish. PLoS Pathog 9, e1003619, doi:10.1371/journal.ppat.1003619 (2013).

18 Wallace, K. N., Akhter, S., Smith, E. M., Lorent, K. & Pack, M. Intestinal growth and differentiation in zebrafish. Mechanisms of Development 122, 157–173, doi:http://dx.doi.org/10.1016/i.mod.2004.10.009 (2005).

19 Gerbe, F., Legraverend, C. & Jay, P. The intestinal epithelium tuft cells: specification and function. Cellular and molecular life sciences : CMLS 69, 2907–2917, doi:10.1007/s00018-012-0984-7 (2012).

20 Karst, S. M., Wobus, C. E., Lay, M., Davidson, J. & Virgin, H. W. t. STAT1-dependent innate immunity to a Norwalk-like virus. Science 299, 1575–1578, doi:10.1126/science.1077905 (2003).

21 Arthur, S. E., Sorgeloos, F., Hosmillo, M. & Goodfellow, I. Epigenetic suppression of interferon lambda receptor expression leads to enhanced HuNoV replication in vitro. bioRxiv, 523282, doi:10.1101/523282 (2019).

22 Kolawole, A. O., Rocha-Pereira, J., Elftman, M. D., Neyts, J. & Wobus, C. E. Inhibition of human norovirus by a viral polymerase inhibitor in the B cell culture system and in the mouse model. Antiviral Res 132, 46–49, doi:10.1016/j.antiviral.2016.05.011 (2016).

23 Rocha-Pereira, J. et al. The viral polymerase inhibitor 2’-C-methylcytidine inhibits Norwalk virus replication and protects against norovirus-induced diarrhea and mortality in a mouse model. J Virol 87, 11798–11805, doi:10.1128/JVI.02064-13 (2013).

24 Legoff, J. et al. The eukaryotic gut virome in hematopoietic stem cell transplantation: new clues in enteric graft-versus-host disease. Nature medicine 23, 1080–1085, doi:10.1038/nm.4380 (2017).

25 Haga, K. et al. Functional receptor molecules CD300lf and CD300ld within the CD300 family enable murine noroviruses to infect cells. Proc Natl Acad Sci U S A 113, E6248–e6255, doi:10.1073/pnas.1605575113 (2016).

26 Murayama, E. et al. Tracing hematopoietic precursor migration to successive hematopoietic organs during zebrafish development. Immunity 25, 963–975, doi:10.1016/j.immuni.2006.10.015 (2006).

27 Karandikar, U. C. et al. Detection of human norovirus in intestinal biopsies from immunocompromised transplant patients. J Gen Virol 97, 2291–2300, doi:10.1099/jgv.0.000545 (2016).

28 Tan, M. & Jiang, X. Norovirus gastroenteritis, carbohydrate receptors, and animal models. PLoS Pathog 6, e1000983, doi:10.1371/journal.ppat.1000983 (2010).

29 Lindesmith, L. et al. Human susceptibility and resistance to Norwalk virus infection. Nature medicine 9, 548–553, doi:10.1038/nm860 (2003).

30 Takamatsu, S. et al. Core-fucosylation plays a pivotal role in hepatitis B pseudo virus infection: a possible implication for HBV glycotherapy. Glycobiology 26, 1180–1189, doi:10.1093/glycob/cww067 (2016).

31 Kettleborough, R. N. et al. A systematic genome-wide analysis of zebrafish protein-coding gene function. Nature 496, 494–497, doi:10.1038/nature11992 (2013).

