## Supplementary data file for "A robust human norovirus replication model in zebrafish larvae"

**Supplementary Materials:**

Figures S1-S7

Tables S1-S4

Materials and Methods

References (31-46)

**SUPPLEMENTARY FIGURES**

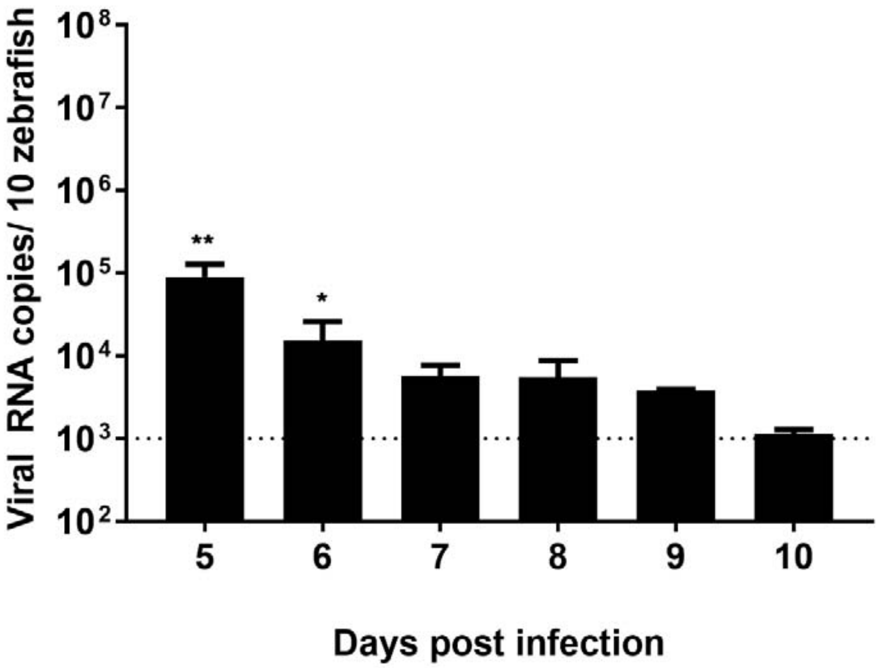


**Fig. S1. HuNoV GII.P7-GII.6 is detected in zebrafish larvae until 6 days pi.** Infection of zebrafish larvae (3dpf) with HuNoV GII.P7-GII.6 (3 independent experiments), zebrafish larvae were harvested each day pi, bars represent the mean values ± SEM of viral RNA levels/10 zebrafish larvae as quantified by RT-qPCR. The dotted line represents the limit of detection (LOD). In every independent experiment 10 zebrafish larvae were harvested at each time point, Mann-Whitney test, where, **p<0.01, *p<0.05.

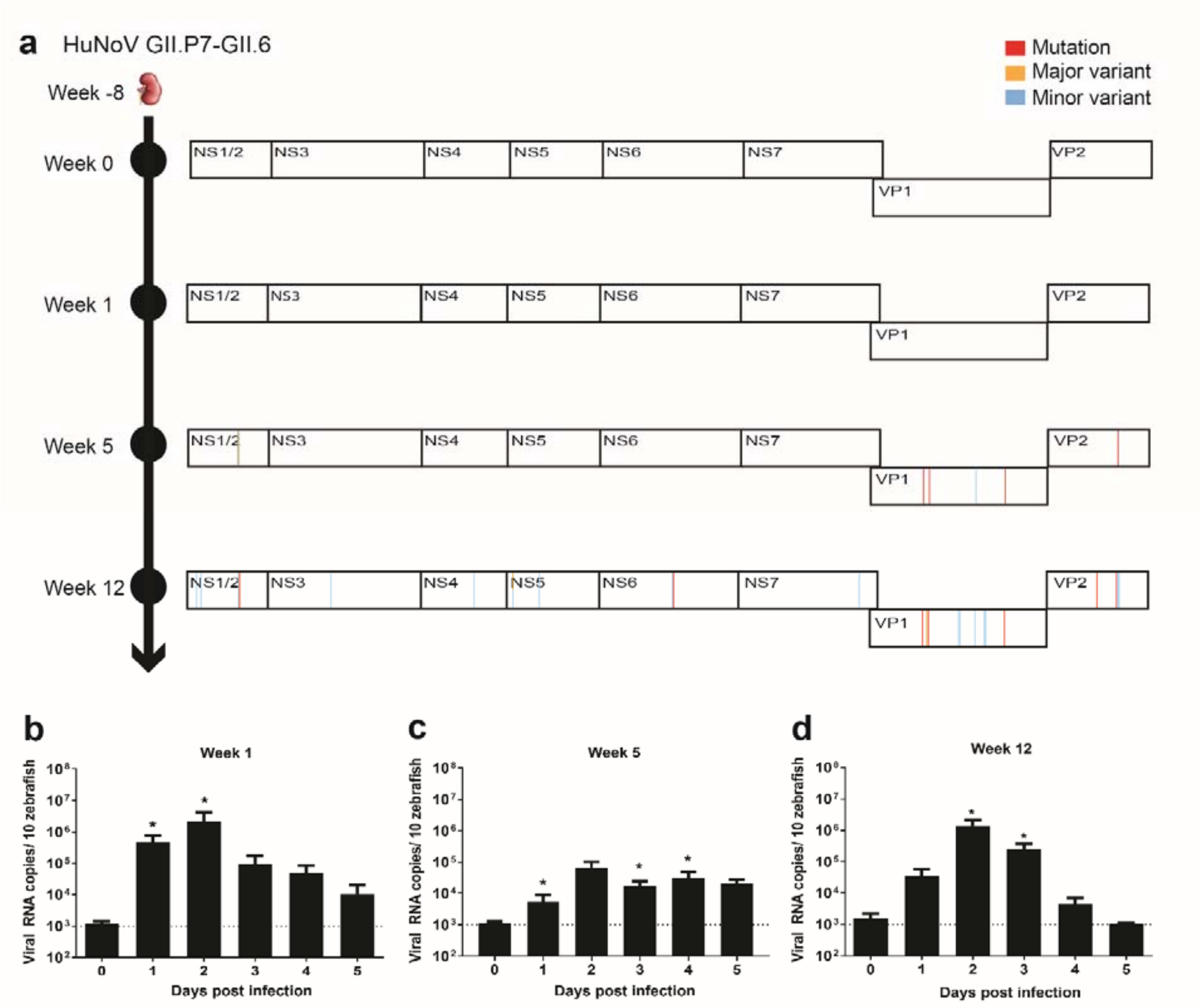


**Fig. S2. Characterization of the HuNoV GII.P7-GII.6 sample and clinical history of the patient.**

(**a**) A 2.5-year-old patient that had received a kidney transplant and presented 8 weeks later with acute viral diarrhea caused by HuNoV GII.P7-GII.6. The infection lasted for ~7 months along which multiple stool samples were collected; four of these samples were characterized (week 0, week 1, week 5 and week 12). The immunosuppressive therapy consisted of tacrolimus and mycophenolate. The complete HuNoV genomes were determined by deep sequencing using the NextSeq500 platform (Illumina) and whole genome sequence analysis was performed in comparison to week 0. Mutations, major and minor variants (respectively, ≥80%, 50-80% and 10-49% of the reads had a different nucleotide) are depicted for the HuNoV strains present in samples from week 1, 5 and 12 pi. (**b-d**) Ten zebrafish larvae infected with HuNoV GII.P7-GII.6 of week 1, 5 or 12 were harvested each day pi (4-5 independent experiments). The calculated inocula (3 nL per zebrafish larvae) were (**b**) $2.2x10^4$ (**c**) $2.5x10^3$ and (**d**) $4.4x10^4$ viral RNA copies. Larvae were harvested each day pi, bars represent the mean values ± SEM of viral RNA levels/10 zebrafish larvae as quantified by RT-qPCR. The dotted line represents the LOD. Mann-Whitney test, where $*p<0.05$.

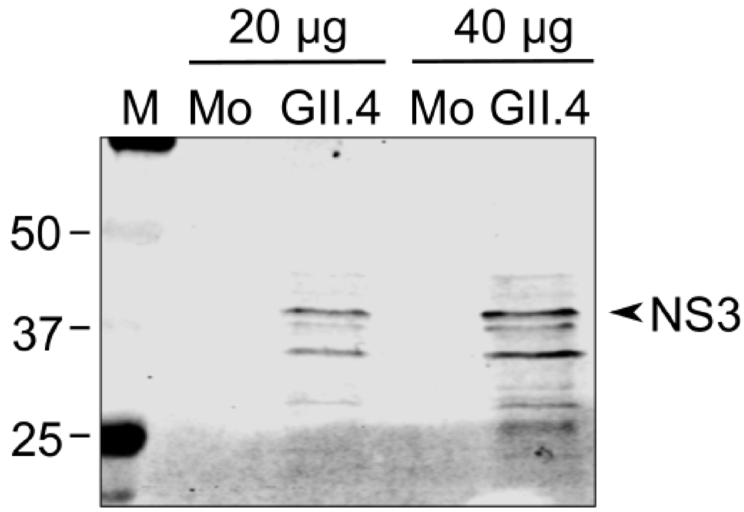


**Fig. S3. Detection of NS3 in HuNoV GII.4 infected zebrafish by western blot.** Western blot analysis of the expression of NS3 in mock (Mo) or HuNoV GII.4-infected zebrafish larvae at 3 days pi. Twenty and 40 µg of the zebrafish larvae lysates were loaded on the gel. M: molecular weight marker.

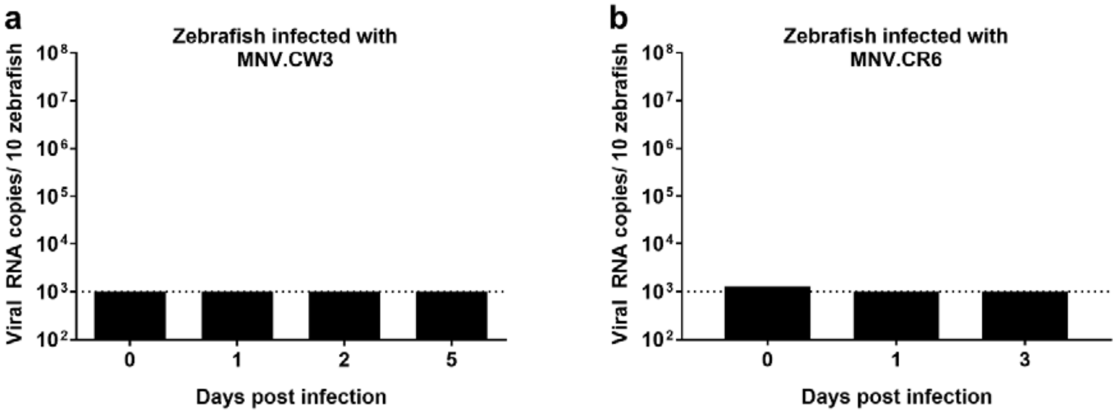


**Fig. S4. MNV.CW3 and MNV.CR6 do not replicate in zebrafish larvae.**
Zebrafish larvae infected with (**a**) MNV.CW3 or (**b**) MNV.CR6 (2-3 independent experiments). Larvae were harvested each day pi, bars represent the mean values ± SEM of viral RNA levels/10 zebrafish larvae as quantified by RT-qPCR. The dotted line represents the LOD.

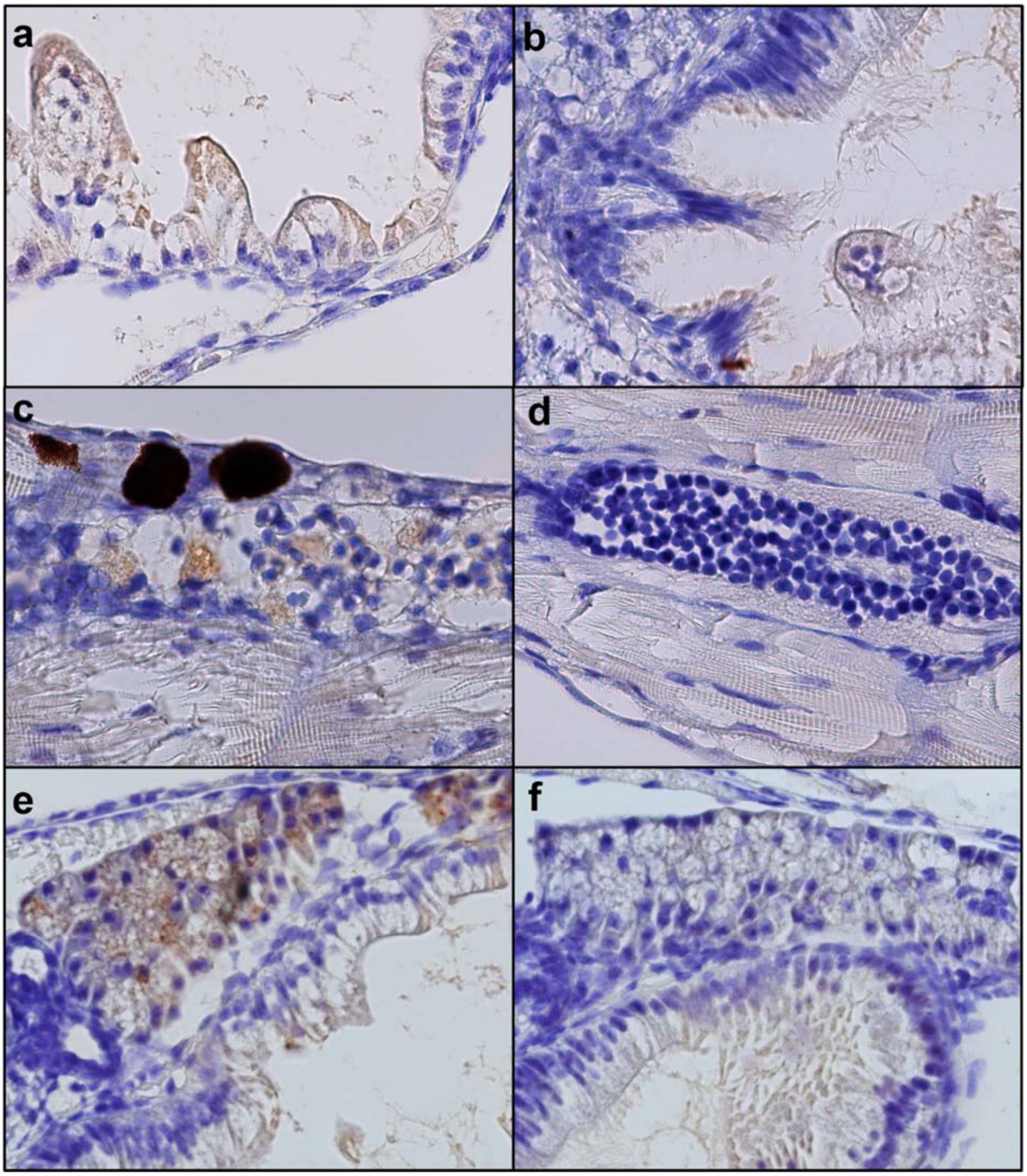


**Fig. S5. Detection of HuNoV VP1 in coronal sections of infected zebrafish larvae.** Immunohistochemistry of coronal sections of HuNoV GII.P7-GII.6-infected larvae (**a,c,e**) harvested at day 3 pi [and the respective uninfected controls (**b,d,f**)]. Images show 5 µm sections stained with VP1-targeting antibodies at 40x magnification, depicting the intestine (**a,b**), CHT (**c,d**) and liver (**e,f**) of infected and uninfected zebrafish larvae.

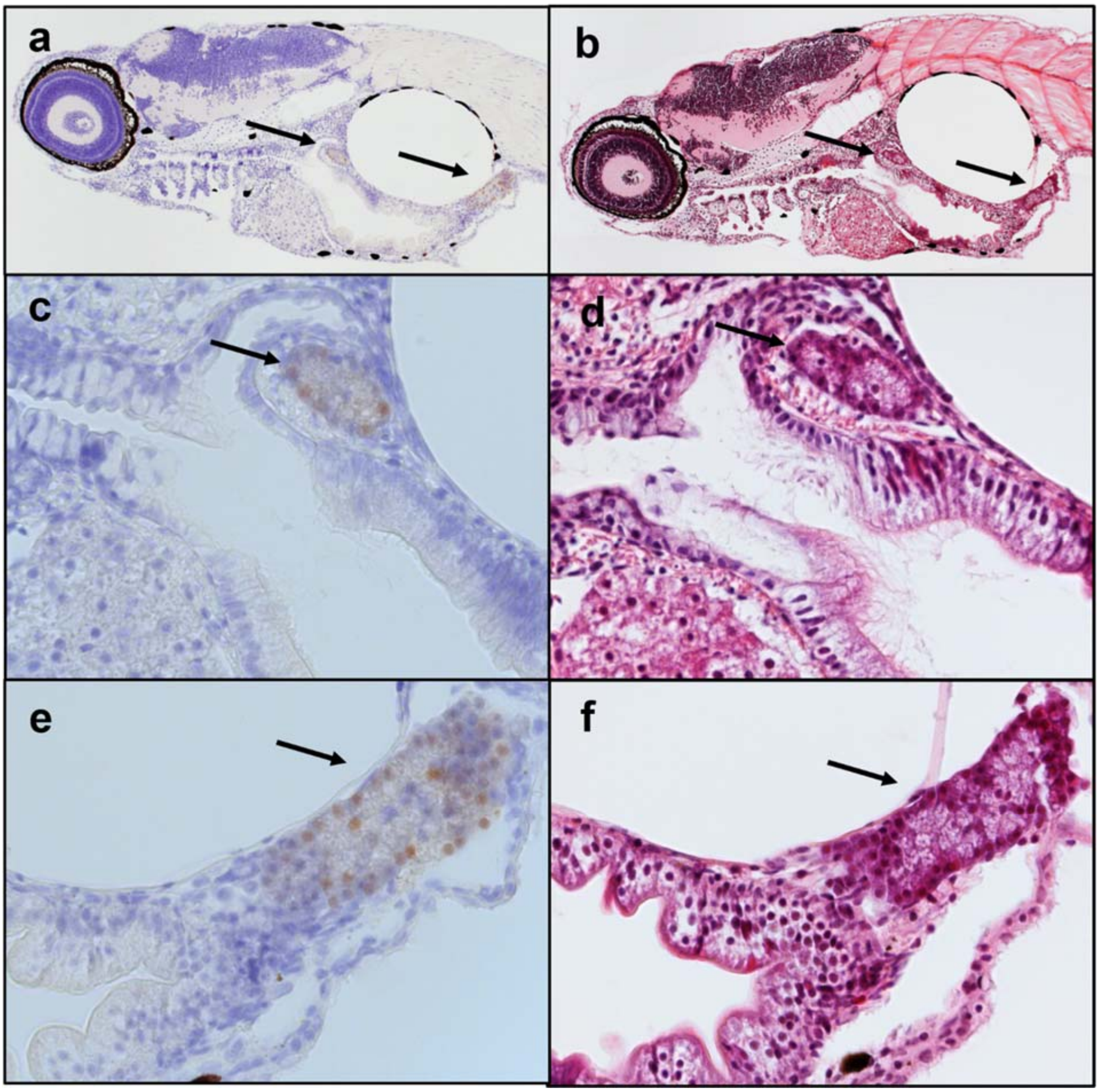


**Fig. S6. Immunohistochemistry and H&E of sagittal sections of infected zebrafish larvae.** Immunohistochemistry with VP1-targeting antibodies (**a,c,e**) and respective H&E staining (**b,d,f**) of sagittal sections of HuNoV GII.P7-GII.6-infected larvae harvested at day 3 pi. Images show 5 µm sections at 10x (**a,b**) and 40x magnifications (**c-f**) highlighting the pancreas (arrows) of infected larvae.

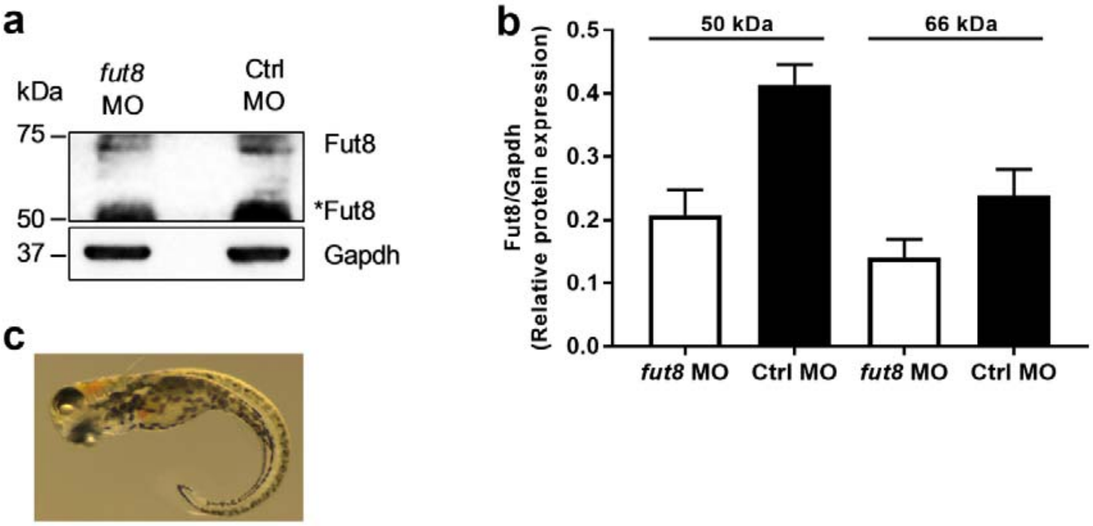


**Fig. S7. Validation of the *fut8* knockdown by western blot.**
Single-cell stage embryos were exposed to a *fut8*-targeting or control morpholino (MO). At 4 dpf, the larvae were harvested for western blot analysis of the expression of Fut8. (**a**) Gapdh was used as a loading control. *Fut8: Probable isoform of Fut8 (**b**) Relative protein expressions, bars represent the mean values ± SEM of 5 independent experiments. (c) A *fut8*-injected zebrafish larvae, presenting with defective midline patterning during development[31].

### SUPPLEMENTARY TABLES:

### Table S1 - Accumulated mutations in HuNoV GII.P7-GII.6 strain over time

| **ORF1** | Week 1 | Week 5 | Week 12 |
|---|---|---|---|
| A82W | - | - | Minor variant |
| T83W | - | - | Minor variant |
| A367R | - | Major variant | Major variant |
| G368R | - | Minor variant | Minor variant |
| A376G | - | - | Mutation |
| A1051R | - | - | Minor variant |
| A2105R | - | - | Minor variant |
| C2392R | - | - | Major variant (A), Minor variant (G) |
| G2585R | - | - | Minor variant |
| A3565R | - | - | Minor variant |
| A3566G | - | - | Mutation |
| A4940R | - | - | Minor variant |
| **ORF2** | Week 1 | Week 5 | Week 12 |
| G576T | - | Mutation | Mutation |
| C617S | - | - | Major variant |
| G630T | - | Mutation | Mutation |
| T910Y | - | - | Minor variant |
| A922R | - | - | Minor variant |
| A1060R | - | Minor variant | Minor variant |
| T1147Y | - | - | Minor variant |
| C1159Y | - | - | Minor variant |
| T1331A | - | Mutation | Mutation |
| **ORF3** | Week 1 | Week 5 | Week 12 |
| Y406C | - | - | Mutation |
| T551Y | Minor variant | - | - |
| T560C | Minor variant | | Mutation |
| T560Y | - | Mutation | - |
| G565R | Minor variant | - | Minor variant |
| C567M | - | - | Minor variant |
| A625R | - | - | Minor variant |

HuNoV GII.P7-GII.6 (week 0) was used as the reference sequence, nucleotide changes in this virus over time (week 1, 5 and 12) were detected. A nucleotide change is defined as a mutation, major or minor variant if respectively ≥ 80%, 50-80% or 10-49% of the reads were different from the reference sequence for a particular nt position.

**Table S2 Overview of *fucosyltransferase* genes present in humans and zebrafish**

| ***Fucosyltransferase* genes present in humans** | **Predicted subcellular localization** | **Protein expression in relevant tissues/cells** | **Orthologues in zebrafish (% similarity, nucleotide)** |
|---|---|---|---|
| ***FUT1***<br>GC19M048748<br>UniProtKB - P19526 | Plasma membrane<br>Golgi apparatus | Colon | NP |
| ***FUT2***<br>GC19P048695<br>UniProtKB - Q10981 | Extracellular<br>Golgi apparatus | Colon (epithelial cells)<br>Ileum (epithelial cells) | NP |
| ***FUT3***<br>GC19M005843<br>UniProtKB - P21217 | Extracellular<br>Golgi apparatus | Colon (epithelial cells)<br>Ileum (epithelial cells) | NP |
| ***FUT4***<br>GC11P094544<br>UniProtKB - P22083 | Golgi apparatus | Colon | NP |
| ***FUT5***<br>GC19M005865<br>UniProtKB - Q11128 | Golgi apparatus | - | NP |
| ***FUT6***<br>GC19M005865<br>UniProtKB - P51993 | Extracellular<br>Golgi apparatus | Ileum (epithelial cells)<br>Colon (epithelial cells) | NP |
| ***FUT7***<br>GC09M137030<br>UniProtKB - Q11130 | Golgi apparatus | - | 53 |
| ***FUT8***<br>GC14P065411<br>UniProtKB - Q9BYC5 | Extracellular<br>Golgi apparatus<br>Cytosol | Ileum (epithelial cells)<br>Colon (epithelial cells)<br>B-lymphocyte<br>Cytotoxic T-lymphocytes | 72 |
| ***FUT9***<br>GC06P096015<br>UniProtKB - Q9Y231 | Golgi apparatus | - | 62 |
| ***FUT10***<br>GC08M033308<br>UniProtKB - Q6P4F1 | Golgi apparatus<br>Nucleus<br>Endoplasmic reticulum | - | 61 |
| ***FUT11***<br>GC10P073772<br>UniProtKB - Q495W5 | Extracellular<br>Golgi apparatus | - | 61 |

Similarity of orthologues is based on nucleotide level. NP: not present.

**Table S3 Primers and probes**

| Name | Sequence 5'-3' | Concentration | Reference |
|---|---|---|---|
| **RT-qPCR** | | | |
| Primers | | | |
| QNIF4 | CGCTGGATGCGNTTCCAT | 900 nM | [32] |
| NV1LCR | CCTTAGACGCCATCATCATTTAC | 900 nM | |
| QNIF2 | ATGTTCAGRTGGATGAGRTTCTCWGA | 500 nM | |
| COG2R | TCGACGCCATCTTCATTCACA | 900 nM | |
| MNV FW | CACGCCACCGATCTGTTCTG | 900 nM | [33] |
| MNV REV | GCGCTGCGCCATCACTC | 900 nM | |
| Probes | | | |
| NVGG1p | FAM/TGGACAGGAGAYCGCRATCT/3BHQ | 250 nM | [32] |
| RING2 | FAM/TGGGAGGGCGATCGCAATCT/3BHQ | 250 nM | [34] |
| MNVprobe | FAM-CGCTTTGGAACAATG-MGB-NFQ | 200 nM | [33] |
| **RT-PCR and genotyping** | | | |
| Primers | | | |
| G1SKF | CTG CCC GAA TTY GTA AAT GA | 1.2 µM | [35] |
| G1SKR | CCA ACC CAR CCA TTR TAC A | 1.2 µM | |
| COG2F | CARGARBCNATGTTYAGRTGGATGAG | 1.2 µM | [34] |
| G2SKR | CCR CCN GCA TRH CCR TTR TAC AT | 1.2 µM | [35] |

**Table S4 Sequencing Coverage for NGS Experiment**

| Week | HuNoV reads | Total trimmed reads |
|---|---|---|
| 0 | 10370 | 3,981,034 |
| 1 | 9823 | 5,004,934 |
| 5 | 2181 | 4,114,059 |
| 12 | 66946 | 5,315,468 |

### MATERIALS AND METHODS

#### Zebrafish maintenance

Wild type AB adult zebrafish were maintained in the aquatic facility of the KU Leuven (temperature of 28 °C and 14/10 h light/dark cycle). Fertilized eggs were collected from adults placed in mating cages and kept in petri dishes containing Danieau's solution (1.5 mM HEPES, 17.4 mM NaCl, 0.21 mM KCl, 0.12 mM $MgSO_4$, and 0.18 mM $Ca(NO_3)_2$ and 0.6 μM methylene blue) at 28 °C until the start of experiments. All zebrafish experiments were performed according to the rules and regulations of the Ethical Committee of KU Leuven (P086/2017).

#### Collection and processing of HuNoV stool samples

Human stool samples, positive for human norovirus (HuNoV), were obtained from the University Hospital of Leuven (Belgium). An aliquot of 100 mg of each stool sample was re-suspended in 1 mL of sterile PBS, thoroughly vortexed and centrifuged (5 min, 1,000 g), and supernatant was harvested and stored at -80 °C. This virus suspension was used for RNA extractions, quantification by RT-qPCR, sequencing and injections in the zebrafish larvae. HuNoV RNA was extracted from 100 μl of PBS suspension using the RNeasy minikit (Qiagen), according to the manufacturer's protocol. UV inactivation of the sample was done by 10 min radiation under an UV lamp (UVP, UVG-54 254 nm). The virus samples used in this study where the following: HuNoV GI.P7-GI.7 ($1.19x10^{12}$ RNA copies/g of stool), HuNoV GII.P16-GII.2 ($7.67x10^{10}$ RNA copies/g of stool), HuNoV GII.P16-GII.3 ($1.58x10^{11}$ RNA copies/g of stool), HuNoV GII.P4-GII.4 ($5.50x10^{12}$ RNA copies/g of stool), HuNoV GII.P4-GII.4 $1.01x10^{12}$ RNA copies/g of stool, HuNoV GII.P17-GII.6 ($1.12x10^{13}$ RNA copies/g of stool), MNV.CW3 ($9.33x10^{6}$ $TCID_{50}$/mL) and MNV.CR6 (9.94 $x10^{8}$ $TCID_{50}$/mL).

#### Infection of zebrafish larvae with HuNoV or MNV by intra-yolk injection

Three dpf zebrafish larvae were anaesthetized by immersion for 2-3 minutes in Danieau's solution containing 0.4 mg/mL tricaine (Sigma-Aldrich, stock solution 4 mg/mL in $Na_2HPO_4$, pH 7-7.5). Thereafter, the zebrafish larvae were transferred to a petri dish (92x16mm) with grooves of a mold imprint (6 rows, one side of 90º the other of 45º) in 1.5% agarose. A dissection needle was used to gently orient the zebrafish larvae so that they were lying on their dorsal side with the yolk facing upwards. Injection needles were pulled using glass capillaries (WPI, TW100F-4) and a Micropipette Puller fitted with a heat filament (Sutter Instruments). In every experiment, the injection needle was calibrated to ensure the precision of the injection volume. Microinjection was done using a M3301R Manual Micromanipulator (WPI) and a Femtojet 4i pressure microinjector (Eppendorf). Each zebrafish larvae was injected with 3 nL of virus (HuNoV GII.2, HuNoV GII.3, HuNoV GII.4, HuNoV GII.6, HuNoV GI.7, MNV.CW3 or MNV.CR6), while negative control zebrafish were injected with 3 nL of PBS. After infection, zebrafish larvae were transferred to 6-well plates with Danieau's solution and further maintained in an incubator with a 14/10 h light/dark cycle at 32 °C. Every day post infection (pi), the general condition of the zebrafish larvae (e.g. posture, swimming behavior or signs of edema) was observed in order to record clinical signs of virus infection, and 10 zebrafish larvae were collected into 2 mL tubes containing 2.8 mm zirconium oxide beads (Precellys/Bertin Technologies) and stored at -80 °C. To determine the 50% infectious dose ($ID_{50}$), 10-fold dilutions up to 1/10,000 of a HuNoV GII.P7-GII.6 PBS suspension were used to infect zebrafish larvae, as described above. The $ID_{50}$ was defined as the virus inoculum necessary to result in the detection of a significant increase of viral RNA lasting for more than one day pi in 50% of infected zebrafish larvae.

**Antiviral treatment**
2′-*C*-methylcytidine (2CMC) was obtained from Carbosynth Limited; a stock solution was prepared in DMSO (VWR Chemicals). Treatment, via immersion, with 4 mM 2CMC started 1 day prior to infection with HuNoV GII.6 or HuNoV GII.4, thus in 2 dpf embryos (10 per condition, manually dechorionated using 2 fine tweezers), and was replenished every 12 h until the end of the experiment. The potential toxicity of 2CMC, was evaluated beforehand and a non-toxic concentration was selected to treat infected zebrafish larvae (data not shown).

**Tissue homogenization and RNA extraction**
Zebrafish larvae harvested in Precellys tubes were homogenized with 3 cycles of 5 sec (6300 rpm) with rest intervals of 30 sec (Precellys24, Bertin Technologies). Homogenates were cleared by centrifugation (5 min, 9,000 g) and RNA was extracted using the RNeasy minikit (Qiagen), according to the manufacturer's protocol.

**RT-qPCR for detection of human and murine norovirus**
For detection of HuNoV GII or MNV RNA, a one-step RT-qPCR was performed using the iTaq Universal Probes One-Step Kit (Bio-Rad), primers and probes used are in Table S3. Cycling conditions were: reverse transcription at 50 °C for 10 min, initial denaturation at 95 °C for 3 min, followed by 40 cycles of amplification (95 °C for 15 s, 60 °C for 30 s) [Roche LightCycler 96, Roche Diagnostics]. For absolute quantification, standard curves were generated using 10-fold dilutions of template DNA of known concentration.

**Sanger sequencing and genotyping**
HuNoV isolated RNA was reverse transcribed by a one-step multiplex RT-PCR using the OneStep RT-PCR Kit (Qiagen), according to the manufacturer's protocol, primers used are in table S3. The cycling conditions were reverse transcription at 50 °C for 30 min, initial denaturation at 95 °C for 15 min, followed by 40 cycles of amplification (94 °C for 30 s, 55 °C for 30 s, 72 °C for 60 s) and final extension of 10 min at 72 °C. The PCR products were run on a 2% agarose gel. All positive PCR samples were purified using ExoSAP-IT (TermoFisher Scientific) and sequenced with the specific GI and GII primer sets. Viral genotypes were determined with the Norovirus Typing Tool Version 2.0 [36].

**Characterization of the immune response by *ifn*, *mx* and *rsad2/viperin* expression following a HuNoV infection**
To generate the cDNA, the ImProm-II™ Reverse Transcription System (Promega) was used. Briefly, a total of 1 µg (ca. 10 µL) of extracted RNA was added to 1 µL of random hexamers and incubated at 70 °C for 5 min, followed by 5 min at 4 °C. To this reaction mix a total volume of 40 µL containing 8 µl of Improm II 5X reaction buffer, 6 mM $MgCl_2$, 0.5 mM deoxynucleoside triphosphate, 40 units of RNase inhibitor, 1 µL of Improm II reverse transcriptase, followed by an incubation at 25 °C for 5 min, 37 °C for 1 h, and 72 °C for 15 min. A qPCR was performed with 4 µL template cDNA using the SsoAdvanced Universal SYBR green supermix, 600 nM of forward and reverse primers for *ifn*, *mx, rsad2/viperin* and the housekeeping genes *β-actin and ef1a*. Primers sequences were as previously described [37,38]. Cycling conditions were: polymerase activation at 95 °C for 3 min followed by 40 cycles of denaturation at 95 °C for 15 s, annealing at 55 °C and extension at 72 °C for 30 sec (Roche LightCycler 96, Roche Diagnostics). Data was normalized to *β-actin* and compared to PBS-injected zebrafish larvae to determine the fold induction of the expression, according to the Livak method [39].

### Sample preparation for viral metagenomics

Human fecal samples were prepared using the NetoVIR protocol, with minor modifications [40]. To preserve the fecal sample for the infection experiments, no virus like particle purification was performed. RNA and DNA were extracted using the QIAamp Viral RNA Mini Kit (Qiagen) according to the manufacturer's instructions, without addition of carrier RNA. First and second strand synthesis and random PCR amplification for 17 cycles were performed using a modified Whole Transcriptome Amplification 2 (WTA2) Kit procedure (Sigma-Aldrich), allowing for amplification of both RNA and DNA [40]. PCR products were purified with MSB Spin PCRapace spin columns (Stratec) as instructed and library preparation was done using a modified Nextera XT DNA kit (Illumina) protocol [40]. Libraries were quantified with the KAPA Library Quantification kit (Kapa Biosystems) and DNA size of libraries was obtained using Agilent High Sensitivity DNA Kit on a Bioanalyzer 2100 (Agilent). Sequencing of the samples was performed on a NextSeq500 platform (Illumina) for 300 cycles (150 bp paired ends).

### Genomic analysis

Raw Illumina reads were trimmed for quality and adapters using Trimmomatic (version 0.35), Table S4. The remaining reads were *de novo* assembled into contigs with SPAdes assembler (version 3.9.0) using the metaspades flag [41]. Contigs were classified using DIAMOND in sensitive mode [42]. A full norovirus genome sequence was obtained for the baseline sample (week 0) which was manually checked by aligning sample reads using BWA-MEM [43]. The assembled human norovirus genome of week 0 was used as reference to align the longitudinal patient samples to infer viral changes over time using BWA-MEM [43] and Tablet [44]. Trimmed reads from each fecal sample were mapped to the baseline sample of the patient (HuNoV GII.P7-GII.6 week 0) using BWA 32 to obtain individual sample magnitudes for the analysis of the donor-derived reads. An in-house developed python script (python version 2.7.6) was used to generate a summary of annotated viral genus reads per individual sample. To assess mutations on the viral genome overtime, samples from week 1, week 5 and week 12 were compared to the sample of week 0 using the package deepSNV from biocLite in R. Sites with less than 10 reads coverage were excluded from the analysis. A site was considered either having a mutation, major or minor variant if respectively ≥ 80%, 50-80% or 10-49% of the reads were different from the reference sequence for a particular nt position. Nucleotide positions with < 100 reads were only included as minor variant if >20% of the reads were mutated compared to the reference.

### Enzyme immunoassay (EIA)

HuNoV structural antigens were detected in infected zebrafish larvae via the RIDASCREEN Norovirus 3rd Generation (R-Biopharm), according to the manufacturer's instructions. Ten (infected) zebrafish larvae were harvested 1 h pi or 3 days pi and deyolked. The zebrafish larvae were smashed in 50 µl of $ddH_2O$ with a pestle in a micro centrifuge tube and debris was removed by centrifugation (10 min, 9,000 g). The supernatant was diluted to a volume of 100 µl. In each EIA run, the positive and negative controls of the kit were included for assay validation and cutoff calculation. The optical density (OD) was measured at 450 nm (Safire, Tecan).

### Gene knockdown by microinjection of morpholino oligonucleotides

Morpholino oligonucleotides (MO) microinjections to target the zebrafish *fucosyltransferase 8* gene were performed in one cell stage embryos with 3 ng of a fluorescein-labelled anti-sense morpholino (5'-CGCCTACTGCTGCCCTCCCCTTTC-3') [31]. As control an equal amount of

GeneTool's standard control morpholino labelled with fluorescein (5'-CCTCTTACCTCAGTTACAATTTATA-3') was used.

**Western blot**

Mock and infected zebrafish larvae were deyolked at 3 days pi, then lysed with a pestle in RIPA buffer (Thermo Scientific) in the presence protease inhibitor (Calbiochem) and centrifuged at 9,000 g for 5 min at 4 °C. Protein concentration of the lysates were determined using the BCA protein assay kit (Thermo Scientific). BHK cells transfected with GII.4 construct were used as positive controls. Thirty µg of either mock or infected zebrafish larvae lysates were analyzed by SDS-PAGE and western blotting. Baby hamster kidney cells expressing T7 polymerase (BSR-T7 cells) were used to analyze viral protein expressions of HuNoV GII.4. Briefly, BSR-T7 cells infected with poxviruses expressing T7 RNA polymerase at an MOI (based on the virus titer in chick embryo fibroblasts) of 0.5–1.0 PFU per cell, were subsequently transfected with 1 µg construct containing the full length clone of HuNoV GII.4 using Lipofectamine 2000 according to the manufacturer's instructions (Invitrogen). To analyze viral protein expression, cells were harvested 24 h post-transfection for western blot analysis. Proteins were separated in 12.5% or 17.5% SDS-PAGE and transferred onto a 0.45 µm nitrocellulose membrane (GVS North America). Membranes were then blocked with 5% milk/PBS-T, washed and incubated overnight with primary antibody against viral proteins NS3 or VPg, which were kindly provided by Professor Ian Goodfellow (University of Cambridge). Membranes were washed extensively, incubated with species-specific secondary antibodies (Li-cor) and viral proteins were detected using Li-cor Odessey CLx imaging system. As housekeeping gene, *β-actin* (Protein Tech 60008-1-Ig) was used. For the confirmation of the knockdown of the *fut8* gene, zebrafish larvae, that had been microinjected with either the *fut8*-targeting or the control morpholino, were deyolked at 24 h pi. Equal amounts of protein extracts (50 µg) were separated by SDS-PAGE (Criterion XT gel, 10% Bis-Tris, Bio-Rad) and transferred onto a PVDF membrane (Trans-Blot Turbo, Bio-Rad). After blocking with SuperBlock (Thermo Scientific) for 1 h, the membranes were incubated overnight with Fut8 antibody (Novus Biologicals, NBP2-58933) at a 1:500 dilution. Membrane was washed extensively and incubated with species-specific HRP-conjugated secondary antibodies (Dako). Proteins were detected using chemiluminescence according to the manufacturer's instructions (Thermo scientific, SuperSignal West Femto Maximum Sensitivity Substrate) and the ChemiDoc MP Imaging System. As housekeeping gene, *gapdh* (Sigma, SAB2701826) was used.

**Infection of zebrafish larvae via immersion**

Four and five dpf zebrafish larvae were immersed in a 1 mL HuNoV GII.P7-GII.6 suspension diluted in Danieau's (containing ~ $10^{11}$ viral RNA copies). After 6 h of exposure to the virus, the larvae were washed with clean Danieau's, and transferred to a well containing new media and further maintained in an incubator with a 14/10 h light/dark cycle at 32 °C. Every day pi, the general condition of the zebrafish larvae (e.g. posture, swimming behavior or signs of edema) was observed in order to record clinical signs of virus infection, and 10 zebrafish larvae were collected into 2 mL tubes containing 2.8 mm zirconium oxide beads (Precellys/Bertin Technologies) and stored at -80 °C.

**Immunohistochemistry**

HuNoV-infected zebrafish larvae were harvested at 3 days pi (i.e. 6 dpf) and fixed in 4% formaldehyde overnight at 4 ºC. The following day the formaldehyde was replaced by 70% ethanol and the zebrafish larvae were embedded in an agarose mold, then processed in paraffin, sectioned and stained as previously described [45]. Immunohistochemistry was performed using antibody TV20 [46] at 1/1000 dilution (kindly provided by Dr. Peter Sander, R-Biopharm) and Anti-VP1 (ab92976, Abcam), 1/500 dilution. Additional staining's were performed with hematoxylin–eosin (H&E). Microscopy was performed using a Carl *Zeiss Axio Imager Z1* microscope at 10, 40 and 100x magnifications and images were captured and processed with the AxioVision 4.8.2.0 software (Zeiss).

**Statistics**

Data was analyzed using GraphPad Prism 7 (Graph-Pad Software) and p values were determined with the nonparametric Mann-Whitney test, where ****p<0.0001, *** p <0.001, ** p <0.01, * p<0.05, and ns is p≥0.05.

REFERENCES


31 Seth, A., Machingo, Q. J., Fritz, A. & Shur, B. D. Core fucosylation is required for midline patterning during zebrafish development. *Dev Dyn* **239**, 3380-3390, doi:10.1002/dvdy.22475 (2010).

32 Stals, A. *et al.* Multiplex real-time RT-PCR for simultaneous detection of GI/GII noroviruses and murine norovirus 1. *J Virol Methods* **161**, 247-253, doi:10.1016/j.jviromet.2009.06.019 (2009).

33 Baert, L. *et al.* Detection of murine norovirus 1 by using plaque assay, transfection assay, and real-time reverse transcription-PCR before and after heat exposure. *Appl Environ Microbiol* **74**, 543-546, doi:10.1128/AEM.01039-07 (2008).

34 Kageyama, T. *et al.* Broadly reactive and highly sensitive assay for Norwalk-like viruses based on real-time quantitative reverse transcription-PCR. *J Clin Microbiol* **41**, 1548-1557 (2003).

35 Kojima, S. *et al.* Genogroup-specific PCR primers for detection of Norwalk-like viruses. *J Virol Methods* **100**, 107-114 (2002).

36 Kroneman, A. *et al.* An automated genotyping tool for enteroviruses and noroviruses. *Journal of clinical virology : the official publication of the Pan American Society for Clinical Virology* **51**, 121-125, doi:10.1016/j.jcv.2011.03.006 (2011).

37 Phelan, P. E. *et al.* Characterization of snakehead rhabdovirus infection in zebrafish (Danio rerio). *J Virol* **79**, 1842-1852, doi:10.1128/jvi.79.3.1842-1852.2005 (2005).

38 Briolat, V. *et al.* Contrasted innate responses to two viruses in zebrafish: insights into the ancestral repertoire of vertebrate IFN-stimulated genes. *J Immunol* **192**, 4328-4341, doi:10.4049/jimmunol.1302611 (2014).

39 Livak, K. J. & Schmittgen, T. D. Analysis of relative gene expression data using real-time quantitative PCR and the 2(-Delta Delta C(T)) Method. *Methods (San Diego, Calif.)* **25**, 402-408, doi:10.1006/meth.2001.1262 (2001).

40 Conceicao-Neto, N. *et al.* Modular approach to customise sample preparation procedures for viral metagenomics: a reproducible protocol for virome analysis. *Scientific reports* **5**, 16532, doi:10.1038/srep16532 (2015).

41 Bolger, A. M., Lohse, M. & Usadel, B. Trimmomatic: a flexible trimmer for Illumina sequence data. *Bioinformatics (Oxford, England)* **30**, 2114-2120, doi:10.1093/bioinformatics/btu170 (2014).

42 Nurk, S. *et al.* Assembling single-cell genomes and mini-metagenomes from chimeric MDA products. *J Comput Biol* **20**, 714-737, doi:10.1089/cmb.2013.0084 (2013).

43 Li, H. *Aligning sequence reads, clone sequences and assembly contigs with BWA-MEM. arXiv Prepr arXiv*, <http://arxiv.org/abs/1303.3997> (2013).

44 Milne, I. *et al.* Using Tablet for visual exploration of second-generation sequencing data. *Briefings in bioinformatics* **14**, 193-202, doi:10.1093/bib/bbs012 (2013).

45 Sabaliauskas, N. A. *et al.* High-throughput zebrafish histology. *Methods (San Diego, Calif.)* **39**, 246-254, doi:10.1016/j.ymeth.2006.03.001 (2006).

46 Parra, G. I. *et al.* Identification of a Broadly Cross-Reactive Epitope in the Inner Shell of the Norovirus Capsid. *PloS one* **8**, e67592, doi:10.1371/journal.pone.0067592 (2013).
